# Ultra-fast cellular contractions in the epithelium of *T. adhaerens* and the “active cohesion” hypothesis

**DOI:** 10.1101/258103

**Authors:** Shahaf Armon, Matthew Storm Bull, Andres Aranda-Diaz, Manu Prakash

## Abstract

By definition of multi-cellularity, all animals need to keep their cells attached and intact, despite internal and external forces. Cohesion between epithelial cells provides this key feature. In order to better understand fundamental limits of this cohesion, we study the epithelium mechanics of an ultra-thin (~25 um) primitive marine animal *Trichoplax adhaerens*, composed essentially of two flat epithelial layers. With no known extra-cellular-matrix and no nerves or muscles, *T. adhaerens* was claimed the “simplest known living animal”, yet is still capable of coordinated locomotion and behavior. Here we report the discovery of the fastest epithelial cellular contractions to date to be found in *T. adhaerens* dorsal epithelium (50% shrinkage of apical cell area within one second, at least an order of magnitude faster than known examples). Live imaging reveals emergent contractile patterns that are mostly sporadic single-cell events, but also include propagating contraction waves across the tissue. We show that cell contraction speed can be explained by current models of non-muscle actin-myosin bundles without load, while the tissue architecture and unique mechanical properties are softening the tissue, minimizing the load on a contracting cell. We propose a hypothesis, in which the physiological role of the contraction dynamics is to avoid tissue rupture (“active cohesion”), a novel concept that can be further applied to engineering of active materials.

**One Sentence Summary:** We report the fastest epithelial cell contractions known to date, show they fit the kinematics arising from current cytoskeletal models, and suggest the extreme tissue dynamics is a means to actively avoid rupture.

## Main Text

Epithelial apical contractions are mostly known to occur during embryonic developmental stages(5–8). These contractions are slow (each contraction lasting minutes to hours) and precisely patterned in both space and time. They play a crucial role in the morphogenesis of the embryo–and then desist. The molecular and mechanical mechanism of contraction in these non-muscle cells, as well as their tissue level control(9–11), are under intensive investigation(*9, 10, 12–17*). Recently, in vitro spreading assays of adult epithelial monolayers showed similarly-slow cellular contractions, though not as canonically patterned(18–23). The triggering and patterning mechanisms of these contractions in somatic tissues are still unknown.

In an evolutionary context, cellular contractions have been suggested to play a role in cohesion and coordination in early animals lacking neurons or muscles. For the sake of coordinated motility, in the lack of cell wall or other inter-cellular structures, contractions were suggested to counter-act and later replace ciliary power(*24–26*). Sponges, a broad class of early divergent animals lacking nerves and muscles, have indeed been reported to use epithelial contractions throughout their adult life, as part of filter feeding, self-cleaning and defense. These contractions are typically in the form of slow peristaltic waves, though quicker “twitch” responses were reported as well (*27–29*). The way in which individual cellular contractions coalesce into contractility patterns and ultimately into behavior in primitive animals is largely unknown. Directly studying “simple”, basal animals provides new perspectives on epithelial function, as well as insights on the evolutionary leap towards multicellularity. Here we study the epithelium of an early divergent marine invertebrate, *Trichoplax adhaerens* as a model “primitive” epithelium. *T. adhaerens* is one of only a handful of animals that lack nerves and muscles (alongside sponges and some parasites(*30*)). As such, it is mostly composed of epithelium (>80% cell count(*31*)). The animal was claimed to be the “simplest” animal known to live today, in metrics like genome length (98Mbp, ~11k genes), count of cell types (6) and body plan (only dorsal-ventral symmetry breaking)(*1, 32*). However, despite its biological minimalism, the animal is capable of coordinated behaviors, like directed locomotion and external digestion(*34*), chemotaxis(*38*), and propagation by fission(*1*). The entire organism is essentially a flat disk (overall ~25um thick), made of two epithelial layers, that is mostly crawling on surfaces. These geometrical facts alone allow easy access for imaging and perturbation. The ventral epithelium consists of columnar cells and is dense with cilia(*33, 34)*, akin to traditional gut epithelium. The ciliary layer is responsible for the animal crawling on substrates.

Very much in contrast, the dorsal epithelial cells are composed of extremely thin confluent tiles (<1um thick) and an overhanging nucleus(*34, 35*) (Fig. 1A). The “T shape” architecture resembles that of sponges’ contractile tissue (exo-pinacoderm)(*36*). Only adherens junctions were found in electron microscopy imaging of both epithelia(*37*) and no extra-cellular matrix or basement membrane have been seen(*1, 30*) (additional details in Supplementary Data 1).

**Figure 1:**
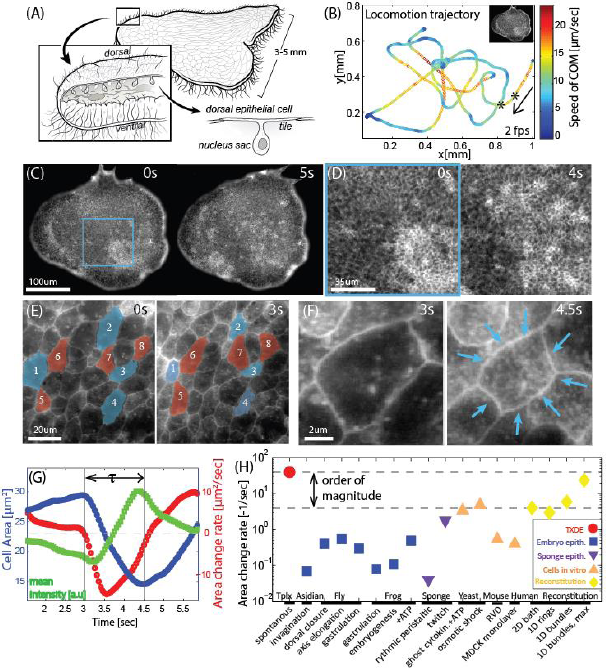
Contractions dynamics in *T. Adhaerens* dorsal epithelium (TADE) at all scales. (A) *T. Adhaerens* consists of two flat cell layers of dorsal and ventral epithelium. The dorsal cell tiles are flat with junctions to neighboring cells (Supplementary Data 1). (B) An example trajectory of an animal’s center-of mass, registered while it is freely crawling in 2D and physically tracked for min. The imaging, tracking and plotting rate is fps. Color represents momentary velocity in the horizontal plane. Animal rotation is not shown. Inset depicts the relative organism size. The segment between the 2 asterisks corresponds to the images in (C). (C-F) Live imaging of TADE across different length scales, from ‘in toto’ to a single cell. Cell Mask Orange (CMO) is used as a live membrane stain at all scales. (C-D) Snapshots from a low magnification movie (Mov. 4): cells with smaller sizes are seen brighter due to increased fluorescent signal. Spatio-temporal patterns are seen in time scales of a few seconds. (E) Snapshots from a high magnification movie (Mov. 5): Contraction events are mostly asynchronous, though correlated to neighboring expansions, as the tissue is always kept intact. Cells labeled blue were contracting and labeled red were expanding in that time interval. (F) A single cell contraction (Mov. 5 top right) (G) Dynamical measures of the single contraction event in (F), after applying computational segmentation to measure the cell’s apical area. The overall area reduction in this event is 50% from initial area, and the contraction duration, *τ*, is 1.5sec. (H) Comparative chart of data from literature reporting epithelium contraction speeds across the animal kingdom and other relevant contraction speeds (citations and comments in Supplementary Data 2).

In this work we use live in-toto imaging and tracking tools to study *T. adhaerens* dorsal epithelium and explore new limits of epithelial contractility and integrity. First we quantitatively describe the contractility phenomena, suggesting the tissue behaves as a highly dynamic active solid. We show that the fast contraction speeds observed are feasible within current models of random cytoskeletal bundles without load. We then bring a list of morphological and physiological evidence to show that the tissue is indeed minimizing the load on a contracting cell. In particular, we demonstrate an extreme dynamic range in cell size and shape, in response to either external or internal forces, making the tissue surrounding a contraction effectively soft. In the discussion we propose that an interplay between contractility and softening could be a means to keep tissue integrity under extensile stress, a mechanism we call “active cohesion”.

### Live ‘in toto’ imaging reveals ultra-fast cellular contractions

We image live animals from top (dorsal) view, as they crawl freely in two dimensions inside a 30um thick micro-fluidic chamber (see methods). By labelling cell membranes with a fluorescent dye (Cell Mask Orange, CMO), cell borders are visualized in the live animal. In order to be able to follow the same cells over long durations in a fast-moving animal, we use an automated 2D tracking system (software controlled microscope stage). To improve cell tracking, we further use post-processing registration algorithms (SIFT, ImageJ) to center the animal and correct for rotations (see methods). Throughout the imaging, animals are crawling on the glass substrate in non-trivial trajectories and speeds of up to um/sec (Fig. 1B), similarly to their behavior without confinement.

Under low magnification, flashes of increased fluorescence are visible in the dorsal epithelium (Mov. 1). Increasing magnification to see individual cells, while maintaining the whole animal in the field of view (‘in toto’ imaging), reveals that the fluorescence dynamics is driven by fast cellular contractions. During these events, fluorophore density in the apical cell surface is increasing (Fig. 1C,D,E,F Mov 2-4). Throughout our data collection, over minutes of imaging, this contraction/expansion dynamics stays active. In addition, the dorsal tissue stayed intact, with no visible ruptures or cellular rearrangements at these time scales.

Following a single cell during an activity cycle demonstrates the ultra-fast kinematics. The cycle includes an oversize expansion phase, followed by a fast, concentric contraction of about 50% in cell apical area within roughly one second, and finally a slower expansion phase, fully recovering the apical area to baseline (Mov. 5, Fig. 1F,G). Quantitative increase in the fluorescent signal tracks the contraction event reliably (Fig. 1G). A literature survey reveals that the contraction speed in *T. adhaerens* dorsal epithelium is at least an order of magnitude faster than any previously recorded epithelial cell contraction. Only recent in-vitro molecular assays of actin-myosin bundles, without any cellular infrastructure or constraints, were shown to reach comparable contraction speeds(*13*) (Fig. 1H, Supplementary Data 2).

In order to extract statistical data of the contractility in *T. Adhaerens* dorsal epithelium, we apply computational segmentation and tracking techniques on to a movie showing sparse contractions (Mov. 6, Fig. 2A,B, methods) and extract the area of individual cells over time. We arbitrarily define an active contraction event as a monotonic decrease in cell area that is longer than 0.8sec. Analyzing all cell tracks, we show that 10-15% of the cells in the monolayer contracted during the one minute movie (Fig. 2B inset). A few examples of the longest trajectories show cells that contracted once, multiple times or not at all. (Fig. 2C). Statistical distributions of all contractile events show (Fig. 2F-I, Extended Data Fig. 1A-B) that on average, a contraction reduces 22um^2^ (or 45%) of cell area during 1.1 second, while peak speeds reach on average 27um^2^/s (or 85%/sec). In cells that exhibit sequential events, the time gap between them was most likely 5 sec (Extended Data Fig. 1C). Averaging the time dynamics of all contraction events reinforces the fast and short contractility acceleration phase with a longer and slower deceleration phase, finally followed by a long cell expansion phase (Fig. 2J-K). We further show that larger cells exhibit higher area reduction (denoted ‘amplitude’, Δ*A*) although the normalized amplitude (Δ*A/A_0_*) is independent of initial cell size. Larger cells also reach higher peak speeds, both in raw and normalized units 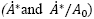. The correlations act as benchmarks for our models.

**Figure 2:**
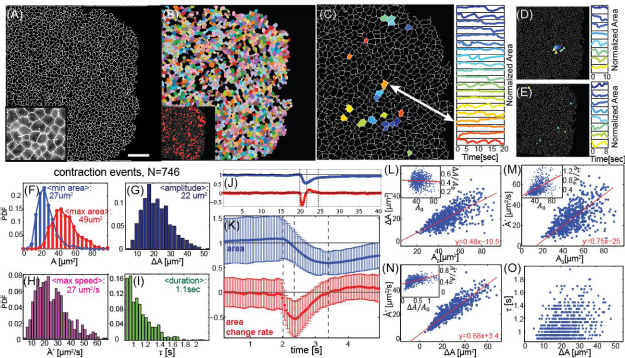
Individual contraction statistics. (A) A single frame from a one minute long movie of a live animal. Computational segmentation identifying cell borders (inset) find roughly 2000 dorsal epithelial cells in a frame (Mov. 6). Scale bar: 50um. (B) Post segmentation, individual cells are marked (color) if they have been tracked for more than two seconds. Inset: Labelled red are all cells that were identified to be contracting sometime during the one minute movie. (C) cells, that were tracked for the longest durations, and their apical area dynamics with time. Color represents cell identity both in the locations map and in the area profiles plot. Area is normalized to be fraction from maximal area. Cells exhibit different behaviors: from no contraction at all, to single events and repeated events (D) 8 neighboring cells and their normalized area dynamics. Color represents cell identity. (E) cells that were undergoing contraction at the same time and their following dynamics. No clear correlation is seen in (D,E). (F-I) Statistical distributions of the 746 contraction events found in all cells trajectories. We plot the initial and final areas (*A*). the amplitude (Δ*A*), peak speed (*A\**) and duration (*τ*) and mention their mean. (J) Aligning all contraction events in time, and normalizing to initial area, we plot the average dynamics of a cell area (*A(t)/A_0_)*, in blue, and area change rate (*A(t) – A(t – 1))/A(t)*, in red. (K) Zoomed in view of (J). Error bars are the variance within the different events in a given time point. (L-O) Correlation plots depict that contraction amplitude and speed are positively correlated to initial cell size and to each other, while duration of the contraction event is independent. Each data point shown is a single contraction event.

### Spatio-temporal contraction patterns across the tissue

Next, we examine the spatiotemporal contraction patterns in the dorsal epithelium. We use particle image velocimetry (PIV) to estimate the local rate of area change by calculating the divergence of the displacement field (see methods). By eliminating low divergence values we highlight the contraction/expansion events (Fig. 3A-C, Mov. 2-4,6, methods). Although the repertoire of patterns is vast, a few simple motifs can be identified. Most commonly seen are sparse, sporadic and asynchronous contractions of individual cells, occurring throughout the tissue (Fig. 3A, Mov. 2,6). Sometimes all cells in the tissue exhibit active cycles, mostly either in or out of phase from each other (Mov. 7). Occasionally traveling waves of contraction are visible, with either radial (Fig. 3B,D Mov. 3) or uniaxial propagation (Fig. 3C,E, Mov. 4). We roughly measure the average wave front propagation in radial waves (0.75 cells/sec, n = 4) to be times slower than that of uniaxial waves (3 cells/sec, n = 4). The cellular signal relevant for this cell-cell interaction is faster than actin turnover. However, the wave fronts are slower than diffusion of small molecules in water and similar to calcium wave propagation through a cell membrane (Fig. 3F). We further notice that wave fronts can both split or merge, and propagate either faster or slower than the gliding speed of the organism itself on the substrate (Mov. 4), implying the tissue acts as a nonlinear excitable medium.

**Figure 3:**
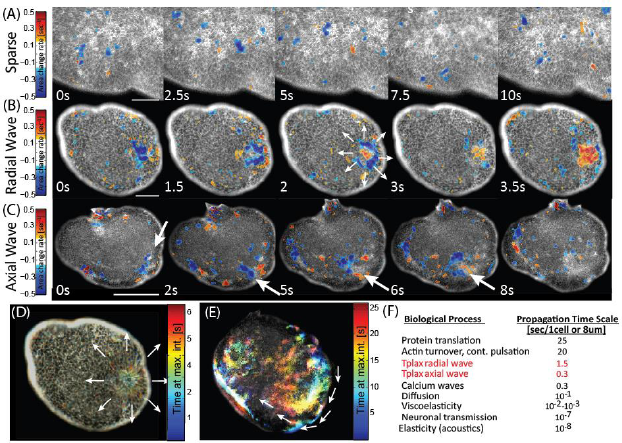
Contractility patterns. (A-C) Examples of common contractile patterns seen in TADE (Mov. 2,3,4). The raw image in underlying a color representation of the divergence field calculated using PIV. Blue range represents negative divergence (i.e. contraction), red range is positive divergence (expansion). Low values of both contractions and expansion are excluded, for clarity. The negative threshold value (−15%area/sec) we define to represent ‘active’ contractions. The positive value is chosen arbitrarily to be the same. Since contractions are faster than expansions, red spots are less common. White arrows mark a propagation of a contraction wave. (A) Sparse contractions of mostly individual cells. Scale bar: 40um. (B) A radially propagating contraction wave that starts at the bulk of the tissue and propagates in all directions. Scale bar: 100um. (C) An axially propagating contraction wave that initially follows the animal rim and then disperses into the bulk. Scale bar: 200um. (D-E) The same events as in (B-C) presented in a color-time technique, similar to ‘maximal intensity projection’ (methods). (F) Characteristic time scales for different cell-cell signaling mechanisms. Citations and comments in Supplementary Data 2.

### The measured high contraction speeds are feasible within actin-myosin bundle speed limits

Actin-myosin networks are known to govern cellular contractions in animal cells, and are suspected to do so also in sponges’ pinacocytes(*27, 39, 40*). We show that F-actin bundles appear as circumferential rings on the apical surface of dorsal cells in fixed, relaxed animals (Extended Data Fig. 2G, methods). This actin ring geometry is similar to other non-muscle contractile cells and is consistent with previous studies of *T. adhaerens* morphology(*31, 37*). We further show that in animals that were not relaxed before fixation, some small cells are located in the center of rosette-like formations, indicating they are frozen in an act of contraction and hence pulling on their neighbors. These cells exhibit high actin density throughout their apical surface (Extended Data Fig. 2H-J).

The high contraction speeds rule out actuation purely by actin assembly or turnover (in which relevant time scales are ~20sec(*41*)) and suggest actuation by myosin motors activity in a given actin geometry. Interestingly, these quick actuation times may be explained by the recently shown criticality in the transition from bundle stability to contractility(*12*). The exact identity of the motors is currently unknown. However homologs of different human myosins (including non-muscle and skeletal myosin II), as well as key non-muscle regulatory factors, were found in the *T. adhaerens* genome using protein sequence alignment (Supplementary Data 3). So far we were unable to significantly inhibit contractile activity using common inhibitory drugs (Blebbistatin, cytochalasin, Latrinculin and ML7, see Supplementary Data 3).

We now perform a calculation to show that current acto-myosin contraction models can explain the high contraction speeds we observe, by bundle amplification alone without counter force. Both the quasi-sarcomeric polarization model and the random polarization model (Supplementary Data 4a) predict similar or even faster speeds than our measurement, using very feasible parameters. That is true even if we assume the actuator to be the common non-muscle-myosin-II, which is the slowest motor known to participate in cellular contractions (and is found in *T. adhaerens* genome, Supplementary Data 3).

We show that a bundle of initial length *L*_0_ that is free in space will shrink with an approximate speed of 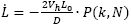 (Fig 4A-B, Supplementary Data 4), where *V_h_* is the speed of a single motor head (as obtained from motility assays) and D is the average longitudinal distance between cross linkers. We introduce *P* as a contraction probability factor: the percentage of motor filaments that will in fact activate contraction. For quasi-sarcomeric geometry *P* = 1. For a randomly polarized actin geometry, we calculate *P* using combinatorics, assuming a buckling-type model and equal probability for all actin orientation micro-states. P in that case is a function of k, the average number of heads in a motor filament that are connected to the bundle, and N, the average number of parallel actin filaments in the bundle (Fig. 4A-C, Supplementary Data 4). For this calculation we allow quick motor attach-detach rates, but assume motors are occupying all available sites at all times. Note, that in this view 1D shortening speed is constant in time (assuming “fixed” actin disordered geometry throughout the 1 sec contraction). However that speed is an average over the bundle length, as areas along the bundle are either contracting in full or half speed, or are totally inactive, as reported experimentally(12).

**Figure 4:**
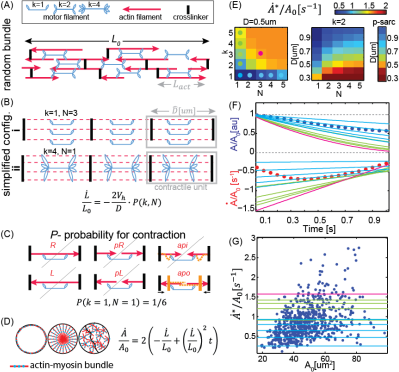
TADE ultra-fast contraction speed can be explained by nm-myosin-II actuation on random actin bundles under minimal load. (A) A randomly polarized acto-myosin bundle at length L_0_. Red arrows represent actin filaments. Arrow points at barbed end. (B) Simplified configurations of random bundles, Using the bundle-averaged parameters: *D*: distance between cross linkers, N: number of actins connected in parallel, and k: number of connected motor head pairs in a single motor filament. Red dotted lines mark actin location axis. Such simplified configurations will yield actin constant shrinking speed 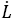 as in the underlying equation. (C) Visualization of all microstates possible for a motor filament in a random bundle with k = N= 1. The microstates are named after the connected actin polarization: Left, Right, Parallel-Left, Parallel-Right, Antiparallel-In, and Antiparallel-Out. According to the buckling model, only one of these states will yield contraction, hence the probability for this unit to contract (denoted by P) is 1/6. (yellow lines represent the state after motor actuation). The combinatorial calculation of P(N,k) is given in Supplementary Data 4. In a quasi-sarcomeric model, P = as only the ‘apo’ state exists (D) Assuming circular cells, all 1D bundles geometries (depicted in dashed red/blue lines) with no load will yield the same area contraction speed 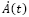 as in the underlying equation. (E) Phase diagrams of the peak area reduction speed 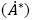 as a function of k, N and D, assuming the buckling model. Colored dots represents parameter sets that we use in the following panels. (F-G) Experimental results (circles) compared to our model predictions for nm-myosin II (lines). Lines color represents the bundle parameters, as depicted in dots on (E). (F) Area and Area change rate as a function of time. Dots are the average data from Fig. 2K. (G) Normalized area change rate as a function of initial cell area. Dots are data from Fig. 2M-inset. The vast majority of the events can be explained by the model in that parameter regime. The few that are higher do not fall within stochasticity (maximal P is 1), but can be explained by either lower D, or measurement errors. The shortening speed predicts complete collapse within ~1sec.

Applying these 1D contractile speeds to 2D circular cell geometry with circumferential bundles (Fig 4D), gives the following dynamics in the apical cell area, 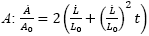. This prediction of linear deceleration in cell area with time suggests the following interpretation to the observed contraction phases: The short acceleration phase corresponds to motor recruitment/activation, while the longer deceleration phase means fully developed, constant bundle contraction speed. Area expansion corresponds to relaxation. Note, that in the lack of load, locating bundles in different geometries in the cell would yield the exact same kinematics, due to the linearity between L and L_0_ (Fig. 4D).

Taking literature value for non-muscle-myosin-II sliding speed without load (*V_h_* = 0.2um/s (*42, 43))*, we can estimate numerical values of bundle contractility rates. A high limit for the contraction speed 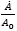 would be 250%area/sec (taking *P* = *1* and *D* = L_myo_ = 0.325um, the length of the motor filament). The low speed limit would be 10%area/sec (taking a random bundle with *k* = *1*, *N* =*1*, and *D* = *2_Lact_+L_myo_* = *1.325um* – the minimal serial network that is still percolated). All our experimental measurements are within this predicted range. We further plot a phase diagram for the area contraction speed as a function of k, N and D in a randomly polarized bundle (Fig. 4E left). Interestingly, the non-linearity of P(k,N) effectively makes P = quite quickly. Hence in the low load regime adding more than a few parallel actins to the bundle, or heads to the motor filament, does not increase the speed significantly (when P = 1 speed values recover the q-sarcomeric model, Fig. 4E right). We plot a few of our model predictions on top of our measured data (Fig. 4F-G). Using *k* = *3*, *N* = *3* (which are low physiological parameters) and *D* = *L_act_* =*0.5um* (as reported in yeast(*44)*) we get speeds that are higher than the average measured contraction dynamics (Fig. 4F), and exceeds 95% of all contraction events observed (Fig. 4G). Further details in Supplementary Data 4.

### Tissue architecture and unique mechanical properties minimize contraction load

How is it, then, that the dorsal epithelium of *T. adhaerens* is the only known example of such fast contractions in epithelia to date? In order to address this question, we turn our attention to morphological and dynamical features that effectively soften the tissue and hence reduce the mechanical load on the motors performing the work. The list of evidence we bring includes in-vivo observations of cell morphology, tissue architecture, membrane morphology, as well as local and global variations in cell size and shape.

Firstly, the confluent part of the tissue is extremely thin: live 3D confocal imaging shows, for the first time in vivo, the unique cell geometry of T-shaped cells, made of ultrathin (≤ 1um) cell pavements while the nuclei are overhanging below (Fig. 5B). Secondly, simultaneous tracking of the dorsal and the ventral epithelia (via splitting a movie by spatial frequencies, see methods) reveals relative horizontal sliding of the two epithelial layers against each other. We measure a relative displacement of up to 70um (~cell diameters) within second (Fig. 5A, Mov. 8). Thus, we deduce at least partial decoupling between the epithelia, as the dorsal layer is essentially suspended on a liquid cavity (Fig. 1A).

**Figure 5:**
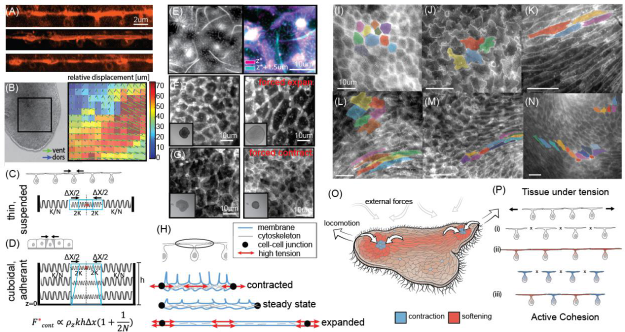
TADE unique and active mechanical properties. (A) A live cross section (XZ plane) of TADE, reconstructed from confocal z stacks. Membranes are labeled with CMO. The cells unique ‘T shape’ is seen, as well as membrane tubes on the apical surface. (B) Left: A snapshot from a movie that shows the animal’s dorsal and ventral epithelia moving independently (Mov. 8). Right: optical plane separation analysis shows relative displacement between epithelia reaching 70um (~10 cells) in 1 sec (C-D) A simplified spring model illustrates that the force required for a single contraction in a thin suspended tissue is lower than in a cuboidal, adherent one (See Supplementary Data 5). (E) A top view of live TADE stained with CMO show membrane tubes. Right: Zoom-in on a single cell, and stacking of the cell borders plane (magenta) and the excursing tubes plane (cyan). (F-G) TADE cells are capable of extreme variation in cell size: (F) Applying compression in the Z direction on the animal results in 200-350% expansion in dorsal cell size before the first visible tear. Insets: whole animal view, FOV: 1.5mm. (G) Treatment with Ionomycine causes immediate contraction of all dorsal cells to approximately 50% within a few seconds (Mov. 9). Inset: whole animal view, FOV: 1.5mm (H) Our hypothetical free body diagrams of a dorsal cell during contraction, expansion and steady states. Red arrows mark regions of high tension and potential rupture in a tissue context (either membrane, cytoskeleton or cell junctions). (I-N) Examples of variable TADE shapes in vivo: polygonal (I), wobbly (J), striated (K), elliptic/amorphic (L, top left). These shapes are commonly found in close proximity in space and change rapidly in time (Mov 10), implying local variability of stiffness. (O) A sketch of the animal from top view, during locomotion. Color represents our hypothetical view of cells increasing (blue) or decreasing (red) their stiffness, and hence their shape, according to different stress patterns. (P) Schematics of possible scenarios of cellular sheets under tension: (i) rupture (ii) expansion due to softening, and then simultaneous contractions and rupture (iii) The Active Cohesion hypothesis suggests active protection against rupture by asynchronous contractions and expansions, activated according to local mechanical cues.

We mathematically formulate the effect of these two features on load reduction using a toy spring model (Fig. 5C-D, Supplementary Data 5): Each cell (of height h) is represented by a parallel stack of horizontal springs of stiffness k at vertical density *ρ_kz_.* N such “cells” are located on each side of the active cell, and the tissue boundary is fixed in place. We compare the force required to achieve the same apical contraction (denoted by Δ*x*) in a thin and suspended tissue, versus in a cuboidal and adherent tissue (such as in embryos). Assuming constant active force producing the contraction, in both suspended and adherent tissues the force is linear with *hΔx* (Fig. 5C,D, Supplementary Data 5). By comparing specific geometries we get that *T.adhaerns* dorsal tissue (h = 0.5um, diameter = 8um) requires a force that is times lower compared to Drosophila embryo right before gastrulation (h = diameter = 8um). This order-of-magnitude calculation shows that the “T-shape” cell geometry significantly reduces the required contractile force to achieve a contraction. Another outcome of the T-shape, with the high area to volume ratio, is the prediction that hydrostatic pressure is not a significant contraction resistance factor, as in spherical/polyhedral cells.

In order to achieve fast contractions, the apical cellular membrane surface also need to match these speeds. We observe a dense collection of membrane tubes protruding from the apical surface of live dorsal epithelial cells (a few dozen tubes per cell, at least 3-4um long. See Fig. 5B,E Extended Data Fig. 2C,F). These membrane tubes were not seen via actin labeling. We hypothesize that these structures are adaptive membrane storage compartments, enabling fast contractions without membrane loss or synthesis, effectively reducing the resistance to expansion/contraction by the membrane. Similar tubular membrane structures have been observed in in-vitro compressed membrane assays(*45*) in which their creation is immediate, passive and purely mechanical.

The capability of vital cells to extremely variate their size is another way for this tissue to reduce contraction resistance forces. To show that, we pursue two perturbation experiments: We compress a live animal in the z-direction, hence apply mostly homogenous, isotropic expansion forces in the XY plane (see methods). In response, the dorsal layer expands immediately to 200-350% of its original area without tearing or any visible damage (Fig. 5F). In comparison, MDCK epithelium cells ruptured at 170-220% expansion of their original size under quasi-static uniaxial stretch over a few minutes(46). In the second experiment, we expose the animal to Ionomycin (50pM in artificial sea water), a drug that increases intracellular Ca2+. The drug triggers contraction of all cells that reduce their area by 50µM within a few seconds (Fig. 5G, Supplementary Data 3, Mov. 9). In both experiments, normal cell size and function are restored after the stress is removed. We conclude, that vital cells can sustain a large range of contracted to expanded states, and quickly move between them. Under physiological conditions, the cells actively maintain an intermediate size (Fig. 5H). After animal’s death, the confluent cells can expand 400%-700% of their steady state size (data not shown).

Finally, live imaging under no externally generated stress shows a wide dynamic range also in individual cell’s apical shape (Fig. 5I-N, Mov. 10). Dorsal cells are usually eliptic/amorphic, however during contraction they become polygonal. Under spontaneous extensile forces (intrinsically driven by the animal’s ventral ciliary layer during locomotion) we notice extremely striated cell shapes. Under spontaneous compression we see wobbly cell edges (shape resembles a jigsaw-puzzle) (Mov 10). Remarkably, these diverse shapes are present right next to each other (Fig 5L). This wide range of shapes infers high, local variability in cortical tension (surface energy), and hence high variability in the effective cells’ stiffness. Cells’ capability to quickly and significantly enlarge/elongate from a base-line size and shape, allows reduction of resistance from neighboring contractions, while keeping the tissue integrity.

## Discussion

A contractile epithelial tissue is an active soft matter system with both continuous and discrete characteristics. Cell cortex, cell junctions and associated cell membrane creates a stress bearing continuum, whereas discrete cells are isolated compartments with finite resources of molecular components, including cytoskeleton filaments, motors and energy sources. Chemical and mechanical thresholds in cell response enable switching behavior between specific cellular mechanical states. Via changing individual cell states, mechanical properties of the entire tissue are affected.

In a contractile epithelium, not only the local geometry, but also the local stiffness is being modulated, especially when considering the “active stiffness” (by this term we mean the integration of intrinsic forces and passive material properties). As an individual cell contracts, it increases its resistance to extensile stresses (i.e. increasing stiffness) both passively (by cytoskeleton connectivity) and actively (by motor activity). However *T.adhaerns* dorsal cells also exhibit dramatic softening, leading to ultra-long striated cells (under tension) or buckles (under compression). Softening can result merely via low cytoskeletal connectivity or low motor activity. The capability of cells and tissues to significantly change their stiffness was suggested before(*47–49*). Specifically stress stiffening followed by stress softening was shown in actin networks in vivo(50). We hypothesize that physiological values of cell size and stiffness in the *T. adhaerens* tissue are actively maintained but can be switched (to stiffer or softer modes) in response to chemical or mechanical cues.

Keeping tissue integrity is at the heart of epithelium function, especially in the case of primitive epithelium. An animal such as *T. adherens*, that is primarily a thin cellular sheet, encounters rapid fluctuations of external forces in the open ocean. In addition, the array of cilia on the ventral epithelium applies locomotion in different directions, which generates physiologically relevant internal stresses (Mov. 10). How does such a seemingly fragile tissue maintain its integrity under these external and internal forces? The capacity to apply retraction forces to counteract extensile stresses is the first step in keeping animal cohesion. However, global simultaneous contraction of all cells under an external constrain will only elevate the tension on cell junctions and potentially cause rupture (Fig 5Pi). Cell softening triggered by an external stretch can reduce tension at cellular junctions, however comprehensive application of that strategy is viable until a maximal expansion limit, after which rupture becomes a threat again (Fig 5Pii).

We hypothesize that *T. adhaerens* tissue integrity is maintained by activation of the two counteracting phenomena - contraction and softening–under different mechanical cues. Quick switching between these two cellular states allows a wide range of responses to immediate stress. We suggest it may allow the application of overall retraction forces, while ascertaining the prevention of critical tensions at cell junctions at all times. We coin this type of mechanism that actively maintains tissue integrity “Active Cohesion” (Fig. 5Piii). The mechanistic details and control of this adaptive mechanism are yet to be studied. Further work is needed also to explore how such multi-state systems can create spatio-temporal contraction/expansion patterns in the presence of physiologically relevant forces. We speculate that similar principles of “active cohesion” may apply in epithelia of higher animals, even if operating at different time scales.

We propose *Trichoplax adhaerens* as a promising model organism for further study of epithelium bio-mechanics, for its many conceptual and experimental advantages. As demonstrated in this current work, studying a broader range of animals beyond the classical model systems may bring completely new perspectives on known biological problems. Our work provides clues for the nature of the putative metazoan ancestor, claimed to be made purely of excitable myoepithlium, the cradle of the creation of nerves and muscles(*25, 26, 51*). Finally, we believe our work will inspire engineering of intrinsically controlled ‘smart materials’ that actively resist rupture.

## Acknowledgements

We thanks Deepak Krishnamurthy, Vivek Prakash, Scott Coyle, William Gilpin, Arjun Bhargava and all Prakash Lab members for discussions and support. We acknowledge Cedric Espenel, Lydia-Marie Joubert and Jon Mulholland for assistance with imaging and Kevin Uhlinger for algae supplies. We thank Chris Lowe for scientific feedback. We finally thank Leo Buss for inspiring discussions, a long friendship, and his generous gift of original animal strains.

## Authors contributions

S.A. performed the experiments and data analysis, with assistance from M.S.B and M.P. A.A.D performed the drugs assays and the protein sequence alignment. S.A and M.P wrote the manuscript. M.P supervised the project.

## Data and materials availability

all data is available in the manuscript or the supplementary materials.

## Supplementary Materials

### Supplementary Materials

available online

### Financial Support

SA was partially supported by Grauss Lipper Foundation and by the Israeli Council for High Education. MP was funded by HHMI Faculty fellowship, Pew Fellowship and NIH Directors Award. This work was partially supported by Chan-Zuckerberg BioHub Investigators Program.

### Author information

Correspondence and requests for materials should be addressed to

## Materials and Methods

### a. Animal culture and sample preparation

Cultures of Trichoplax Adharense from the original Grell strain (1971, courtesy prof Leo Buss) are maintained following a published protocol(*65*). Briefly, organisms are kept in glass petri dishes filled with artificial seawater (ASW. Kent reef salt mix in ddw, at 3% salinity) in 19°c under 18 hours of light conditions per day. Plates are initially seeded with red algae type R-lens (courtesy Kevin Uhlinger from Chris Lowe’s lab at Hopkins marine station) and nutrients (Florida aqua farms 1/4000 volume). 1/3 of the plate water volume is being replaced weekly, maintaining the same nutrient concentration. Live algae is added upon need. Re-plating may be required after 1-2 months.

Prior to any imaging, animals are gently peeled from the plate floor by pipetting and transferred to clean ASW. The shear flows dismiss the debris from the anima’s dorsal surface. Keeping animals in starvation condition for up to 14h further eliminates auto-fluorescence from dorsal and ventral layers.

Imaging chamber is prepared by placing strips of double sided tape (Nilto Denko Corporation, 30um or 10um thick) on a glass slide (24*30mm, #1.5). An animal is put in the chamber inside a known volume droplet (typically 20ul), and let to adhere its ventral side to the slide (5-10min). Closing with a cover slip (22*22mm), the animal is flattened in a known height chamber with a known dorsal-ventral orientation, however free to glide laterally for many millimeters.

### b. Fixation and related staining and imaging

For fixation, staining and drugs application we use a technique of liquids replacement by flow in the chamber itself: after closing a chamber with fluid A by placing a coverslip on top, the chamber is still open on two opposite sides by 30um thick cracks. We place a drop of liquid B on the glass slide next to an ‘open’ side of the chamber and “pull” liquid A from the opposite side, either by using a piece of tissue (for a fast flow in a thick chamber) or by placing a bigger, hence lower curvature, droplet of liquid A on that opposite side (for a slow flow in a thin chamber, using Laplace pressure effect).

#### Fixation

Using the liquid replacement technique, samples are fixed by flowing Lavdowsky solution [50% Ethanol, 10% formaldehyde, 1% acetic acid, 0.1% Tween-20 (*66*)] at −20°c on top of ice, and left at −20°c overnight. Then, washed in PBS three times for 15 minutes each.

#### For SEM imaging

after fixation, the liquid is replaced by a mix of ethanol in PBS in increasing concentrations: 50%, 70%, 90%, 100%, 15 min each. Finally, the liquid is replaced by HMDS in Ethanol (50%, 100%, 15 min each). We peel the top cover slip and allow it to dry in a vacuum desiccator overnight. Samples are then coated with ~10nm gold (Denton Desk II sputter coater). Imaging was performed in in a VPSEM (Hitachi S-3400N).

#### For f-actin staining

prior to fixation, some samples were relaxed by introducing 0.3M sucrose in ASW for 10min(12). After fixation and washing, samples were permeabilized using PBST (PBS+ 0.1%Tween 20, 3 washes of 15min each), blocked in 1% BSA in PBST for hour, stained with Phalloidin (Alexa-Fluor 647 Cell Signaling Technology, 1:40in PBST) for 3 hours and washed with PBST (3 times, 15min each). Images were taken in Zeiss LSM 780 confocal microscope.

### c. Live Imaging

For live fluorescent imaging of cells membrane, we add to the chamber Cell Mask Orange (CMO) 1:500 in ASW (‘Thermo Fisher’). We image the dorsal surface using TE2000-U microscope (Nikon), with lambda XL (Sutter instruments) light source and Orca 4.0 scientific CMOS camera (Hamamatsu). Using a 40x oil objective we achieve ‘in toto’ imaging of ~2000 cells in an animal allowing a single cell resolution. 60x and 100x oil objectives were used as well for higher magnification. Typical movies are taken at 20 fps.

### d. Confocal imaging with physical tracking and z-stacks

For physical tracking of the organism over longer periods of time, we use the Nikon TiE inverted microscope with Andor Neo sCMOS camera, NIS-Elements software and Nikon HCS JOBS software patch. The patch allows stage control according to real-time calculations on each image. We use a 2D cross correlation method to keep the organism in the field of view at all times. Tracked movies are typically taken at 3 fps, for 10 minutes.

We use this optical setting to track an animal at the cell tiles plane, but also to take z stacks that are later reconstructed and presented as XZ cross-section (ImageJ).

### e. Computational registration, cells segmentation and tracking

Acquired movies of 20fps are taken to ImageJ, to enhance contrast and eliminate translations and rotations computationally (SIFT algorithm). Frangi filter algorithm is applied on each frame (Matlab2014), to find lines in the image using the Hessian operator. Finally iterative water-shedding algorithm is used to further segment too large areas at their narrowest point (Matlab2014).

After segmentation is complete on all frames, we label and track individual cells in time by finding minimal cell center displacement (Matlab2014). We allow cells to ‘disappear’ for a few frames while keeping their identity, but we keep only tracks that are longer than two measured seconds.

Area data in time is measured directly. The normalized amplitude of a contraction event is taken to be 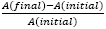. The normalized peak speed is calculated as 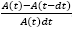. Data is smoothed by a Total Variation Regularization algorithm.

### f. PIV and the color-time technique

Movies undergo PIV (matpiv, Matlab 2014), using interrogation window of 25 um (32 pixels), and time interval of 0.3 seconds. The divergence of the displacement fields is shown, eliminating low absolute values (under 15%/sec). The algorithm uses different objects as “tracers” – from cell walls to subcellular puncta on the membrane.

In parallel, the same movies were processed in a color-time technique, to show time progression in a single image: each frame’s grey levels were translated to intensity levels of a specific color, representing the time point it was taken at. We show the normalized sum of all colored frames. Pixels with distinct color have changed during the movie, and had maximal intensity at the time represented by the color (similar to “maximal intensity projection” representation). Pixels who did not change intensity during the movie, appear in grey level. We present the color map on top of the first frame of the movie.

### g. Spatial frequency split

A movie was taken with Bright field illumination (Nikon Multizoom AZ100) at 20 fps. The animal’s dorsal layer in that setting is mostly transparent however some small features and speckles on it are seen very sharply as they are in mid focal plane. The ventral layer is seen dark in the background. We separate the data from the two optical planes by its spatial frequency (highpass>6% and lowpass<1% of the Fourier transform dimensions). Performing PIV on each of the resulting movies separately, and taking the difference between the displacement fields, gives the relative displacements between the layers.

### h. tissue z-compression

While a live animal is visualized, we apply gentle compression forces in the z direction, either by dropping a flat weight of 35g on the animal in an open petri dish (constant stress setting, Fig 5E low magnification images), or by placing a coverslip on top of the animal that is in a controlled volume droplet, to create an controlled thickness chamber (constant strain setting, Fig 5E high magnification images). Removing the weight in the first case, or adding ASW to the chamber, allows the animal to recover completely and go back to normal cells size.

## Supplementary Text 1: Introduction to *Trichoplax adhaerens*

*Trichoplax adhaerens* is a few millimeters sized, disk-shaped marine animal, found in coastal waters around the world(*1*). It is the only named member of the phylum Placozoa, although world-wide genomic variability has been recently found(*2*). Its exact location in the tree of life is still debated, but most recently it was placed as a sister to Cnideria and Bilateria, later only to Porifera and Ctenaphora(*3-5*). Its genome is small – 98Mbp and approximately 11,500 genes, 87% of them have homologs in other Metazoa(*2*).

Placozoans were claimed to be the simplest of all non-parasitic animals(*6,7*) also due to having only 6 known cell types, and an extremely simple body plan: the animal has no internal digestive lumen (no inside vs outside) and is radially symmetric (no head vs tail). Despite its simplicity, it is capable of complex behaviors, like coordinated external digestion(*8*), probably lead by chemotaxis(*9*), phototaxis(*2*), and propagation by fission(*2*).

Previous work, based on fixed samples, has shown the overall organism structure(*6*) (see also Fig 1A): The dorsal (top) epithelium is made entirely of T shaped cells, exhibiting extremely thin tiles (0.5-1um thick in TEM) and nucleus sacs hanging from them towards the organism interior. We further show a cross section of the tissue in vivo, confirming the live cell morphology (Fig 5b) and show that cell tiles somewhat overlap, hence increase contact area (Extended data Fig 2B). The ventral (bottom) layer is known to be made mostly of columnar epithelium, and their dense, projectile ciliary array is responsible for the animal locomotion by gliding on substrates(*8*).

Between the epithelia is a network of star-shaped fiber cells, with processes touching each other and other cell types (Fig 1A). It was suggested that their inter-connections are synapse-like(*7*) or that the network is a synctium(*10*) but the nature of these connections is still unclear. It was also suggested that fiber-cell contractility governs the overall animal shape changes(*10-12*). however there is no evidence to such contractility to date. Recently a chemical signaling was discovered, related to another cell type: a trigger wave of secretion of endomorphine-like peptide from gland cells, secretory cells in the periphery of the ventral epithelium, governs ciliary beat arrest during feeding. The animal “freezes” in place and “seals” a space underneath it, to allow external digestion and absorption of nutrients. The process takes minutes, and the chemical signal diffuses also outside of the animal(*9*).

The T shaped cells of *T. adhaerens* dorsal epithelium (TADE) resembles that of sponges’ pinacoderm(*13, 14)*, which is also a contractile tissue. In most related sponge species, these interfacial cells contract as part of their filter feeding, self-cleaning and defense(*15, 16*). These contractions are known for their typical peristaltic waves (contraction cycle takes 0.5-3 hours, wave front speed reaching up to 120um/sec), though smaller and faster scale ripples and twitches have also been seen(*15, 17, 18*). Cnideria is another early phylum but with higher complexity, that exhibit both nerves and muscles, as well as myoepithelium (muscles with epithelial features). Interestingly, the myoepithelium shape is similar to TADE cells, only upside down: the striated muscular part is found at the basal side, underlying an epithelial sheet of cuboidal cell bodies. During development, Cnideria cells morph and differentiate from epithelium to either myoepithelium or pure mesenchymal muscle cells(*19*). Hence, this upside down T shape was suggested as a milestone in the evolution of nerves and muscles(*20-23*). These unique cell geometries inspire theoretical thinking about the early animals ancestor made of a contractile epithelial monolayer capable of coordination(*22-26*). Contractility data at the cellular level from both sponges and Cnideria is lacking.

*T. adhaerens* is a fascinating ‘living fossil’, a possible glimpse to events in the evolution of multicellularity, but it also has vast experimental advantages: Its 2D nature makes it accessible for cell-level simultaneous live imaging (‘in toto’) and its biological minimalism allows modeling as a thin sheet of active matter. As demonstrated in this current work, *T. adhaerens* as an emerging model organism may bring new perspectives to existing biological questions.

## Supplementary Text 2: References for speed surveys data (Fig 1H and Fig 3F)

Non muscle cell contraction speeds data (Fig. 1H) is taken from (*27-42*). The numbers are either explicitly mentioned in the papers or estimated from raw data, and adjusted to estimate a contraction speed of 2D circular apical surfaces by a ring-shaped peripheral bundle.

Time scales for cellular and intra-cellular events (Fig 3F) are taken from(*43-48*) except that of viscoelasticity, that was calculated as the ratio between the viscosity of the lipid membrane(*49*) and the young’s modulus of actin filaments(*50*).

## Supplementary Text 3: Identification of cytoskeletal and regulatory molecular components

We looked for potential molecular players by running protein sequence alignment (BLASTp(*51*)) of human genes involved in cytoskeletal contraction to the *T. adhaerens* genome. First, we looked for homologs of muscle and non-muscle myosin II, the myosin type responsible for contractions in other metazoa. We found *T. adhaerens* predicted protein with Uniprot accession code B3RQ46 to resemble human cardiac and type I skeletal muscle myosin II heavy chain, MYH7 (56% identity) and protein B3SCL6 resembles human non-muscle myosin II heavy chain MYH9 (69% identity). We note that current annotation of these genes in the *T. adhaerens* genome includes the myosin motor head but lacks the tail. Furthermore, we found homologs to human non muscle myosin II regulatory light chain MYL9 (75% identity to B3SB79), as well as an essential light chain (MYL6, 56% identity to B3S9B3).

Additionally, we found homologs to human regulatory factors activating smooth and non-muscle myosins. These include calcium-responsive calmodulin (CALM1, 96% identity to B3RJX8) and its target MLC kinase (MYLK, 38% to B3RRX3), Cdc42-binding kinase (MRCKA, 62% to B3SCB2), and Rho (RHOA, 88% to B3RIM6) and its target kinase Citron (CIT, 44% to B3S1X0). No homologs of Rho-inducible ROCK kinase were identified. Homologs to skeletal myosin regulatory troponin complex were not found either. These results are consistent with a previous comparative study of myosin contractile machinery across animal evolution(*52*).

From our homology results, we hypothesized that contractions in TADE cells is regulated by MLCK via phosphorylation of the regulatory light chain. To test this hypothesis we used Ionomycin to increase intracellular Ca^2+^ concentration and activate MYLK, possibly via calmodulin. Indeed, Ionomycin induced instantaneous contraction of all TADE cells (Mov. 9). Taken together, these results suggest that contractions in TADE cells can be elicited by myosin II-like machinery and regulated by canonical factors. Further molecular characterization will be needed to determine whether these proteins have conserved functions and whether this network underlies spontaneous contractions. Transcriptomic characterization of TXDE cells to identify expressed genes in these cells is in progress.

The presence and distribution of f-actin in the cell tiles is discussed in the main text and shown in extended data Fig. 2G-J. The circumferential bundle geometry found is characteristic of non-muscle cells. In line with this, many components of striated muscle Z-discs are missing in the *T. adhaerens* genome, as previously noted(52).

Live cytoskeletal stains (SiR-Actin and SiR tubulin, Spirochrome), as well as inhibitory drugs (Blebbistatin, Cytochalasin, Latrunculin and ML7) failed to stain or to have an effect on spontaneous contractions in TADE cells. This could due to low permeability to these compounds or by divergence in protein sequence and structure, as in the case of Drosophila insensitivity to Blebbistatin(*53*).

Reported motion speeds for active, unloaded non-muscle myosin II are around 0.2 µm/s(*54, 55*). Muscle myosins can be classified as slow (with reported motion speeds ranging from 0.1 to 0.7 µm/s(*56-58*)) or fast (with reported motion speeds ranging from 2 to 8.8 µm/s(*56-59*)). Extensive characterization of myosins expressed in TADE cells will be needed to determine the speed of the motors responsible for their contractions. Nevertheless, while the molecular players are uncertain, our theoretical framework can be used with different molecular parameters, and it essentially shows that a slow candidate for the actuating motor can be amplified in a bundle geometry with minimal load to reach the contraction speeds we see in TADE.

## Supplementary Text 4: Calculating motor speed amplification in 1D acto-myosin bundles

### a) Model-dependent bundle contraction speed

In the last decade, several models have been suggested to explain the ability of acto-myosin bundles to produce net contraction(*60*). In some cases cells adopt a quasi-sarcomeric actin order, in which cross linkers connect the actin’s barbed ends, exposing motors to pointed ends only(*61, 62*). In that geometry, all motors produce contraction. In other contracting cells, bundles are known to have mixed/random actin polarity, and hence the symmetry breaking mechanism is unclear. A leading group of hypotheses is suggesting the idea of a non-linear response of the network to stress, where actin under compression buckles or breaks and hence doesn’t allow force transmission (unlike the case of actin under tension)(*63, 64*).

Here we present a minimal model to calculate fundamental limits of acto-myosin contraction speed in 1D bundles geometry and without load. We consider a bundle with initial length *L*_0_, composed of randomly placed and polarized actin filaments at average length 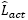, inter-connected by crosslinkers at average longitudinal distance 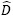 (Fig. 4A). We denote k as the average number of motor-head-pairs actively connected to the actin bundle, N as the average number of parallel motors at play, and we assume the bundle is free in space. In order to estimate the contraction speed, we consider a reduced configuration, composed of 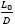 serial contractile units, each occupying N motor filament with k connected head pairs (Fig. 4C). This configuration will shrink with speed 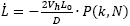 is the speed of a single motor head (as obtained from motility assays). We introduce *P* as a contraction probability factor: the percentage of contraction units that will activate contraction. For a quasi-sarcomeric geometry *P* = 1. For a randomly polarized actin geometry, we will calculate P stochastically as a function of N and k, assuming the buckling model.

### b) Randomly oriented bundle – general model assumptions

In the case of a randomly polarized bundle, we account for all possible actin-orientation microstates. We then assume equal probability for a motor filament to attach to any of these microstates. Estimating ~100 contractile units along a bundle, and averaging over many cells, we can assume that the average of our TADE measurements is a result of sampling all microstates possible many times. We claim local ergodicity and assume the probability of a motor to yield contraction, P, is also the percentage of motors along the bundle that will indeed yield contraction.

Taking the reduced, averaged configuration, we assume homogeneity of motors that occupy 100% of the contractile units. Note, that more than one motor filament connecting the same actins would not contribute to the speed (it may add to the contractile force, which is irrelevant to our no-load model).

We take no assumptions on the duty cycle of the motor filaments, they can detach/attach in new locations in any frequency, as long as the actin geometry is quenched, keeping its statistical properties (which we believe to be reasonable within one second).

### c) Calculation of the contraction probability factor P(k,N)

We now calculate P(k,N) as the probability of N parallel motor filaments, each with k connected head pairs, to induce contraction on a contractile unit. For simplicity, we assume the motor filament is symmetric, i.e. the number of connected heads on each side of the filament is the same.

We label all possible states of a serial heads pair: When such a pair is connected to a single actin, we denote this micro-state as *L* (left) or *R* (right) (direction is pointing towards a barbed end). A state in which a heads pair connects two parallel actins will be marked *pL* or *pR.* The state of connecting anti-parallel actins will be marked *api* (‘in’) or *apo* (‘out’) (Fig4 B). Hence, each pair within the motor filament gets one sample from a space of 6 {L,R,pL,pR,api,apo}. For example: a motor with 4 heads (k = 2) can be in a state *{L-apo}* or *{api-pR}* etc, a 6 headed motor (k = 3) may be found in states like *{pL-L-apo}, {api-api-R}* etc. Therefore, the number of possible states for a motor filament is calculated as how many different *k* samples could be taken from a pool of 6, with possible repetitions and regardless of order. The number of states is given by:

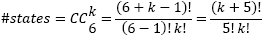

We explicitly depict all possible states for k = 1 in Fig 4B and for k = 2 in Extended Data Fig 3.

We note that a motor filament will not yield contraction when all actins it connects are oriented in the same orientation (i.e. when all k samples are either ‘*L*’ or ‘*pL*’ and when all samples are either *‘R’* or ‘*pR*’). Another case of no contraction at all is when all barbed ends are pointed inwards (i.e. all k samples are ‘*api*’), which is one state only. We take this case to be static under the assumptions of a buckling model. The number of non-contracting states is then:

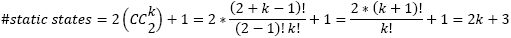

Some microstates are contracting at Vh, which is half of the maximal speed. These are all cases where only one motor head is pulling inwards. That happens in the absence of *‘apo’.* with at least one ‘*api’* and all the rest are pointing in the same direction. That is:

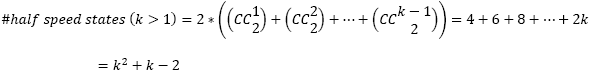

The states of full contraction speed are those where at least one pair is antiparallel outwards. It is either with at least one ‘apo’, or with at least two anti-parallel branches. However it is easier to calculate it as the number of remaining states:

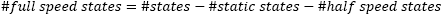

The factor P is the generalized probability, i.e. the weighted average of the speed:

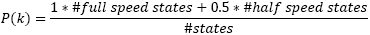

Calculating a few examples:

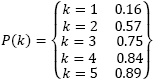

Hence, for example for a four-headed motor (k = 2) there is a 57% chance for contraction. More connected heads (k = 3,4,5..) the probability is rising highly non linearly.

Similar effect happens when adding more k-headed motors in parallel to a contractile unit, only this time every additional motor has the same fixed probability to yield contraction, P(k). The probability for at least one success in a collection of independent trials, when the probability of success in each is P(k), is given by the binomial theorem:

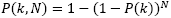

Calculating a few examples:

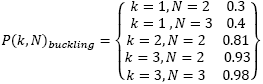

### Supplementary Text 5: Load minimization via cellular and tissue architecture

We bring a reduced model of cell contraction in TADE, as an active spring contracting in the bulk of a passive elastic network with fixed boundaries far away. For simplicity we consider such network in 2D (an XY cross section, see Fig 5C-D). Each cell is represented as a stack of N horizontal springs of stiffness k on each side of the active one. For simplification, we take the central spring midpoint as fixed in space. In this frame of reference, we treat the active spring as made of two serial springs of stiffness 2k each. The rest of the springs can be effectively represented as a single spring with stiffness k/N on each side (Fig 5C-D). To achieve an overall compression of displacement Δ*x* (Δx/2 on each side) the work of the active spring W, or the stored potential energy Ep, will be:

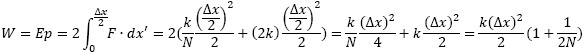

In the calculation we included both the energy stored in the compression of the active spring and the one stored in the tension of the rest of the network. In a finite tissue thickness h, and spring density of *ρ_kz_* we get a ‘stiffening’ effect, and the stored energy will be linear with h:

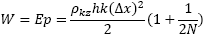

Assuming constant active force acting throughout the contraction, its value would be:

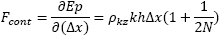

Note, that the bigger the organism (higher N), the softer (lower k), and the thinner are the cells (lower h) - the lower is the force required for contraction at an amplitude Δ*x*.

For comparison, a contraction of a cuboidal cell in a flat tissue with only its apical surface being active, while the basal side is adherent to a bottom substrate, produces a trapezoid cross section. Due to horizontal springs we get:

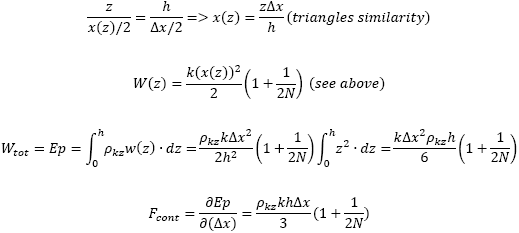

In both cases the force is linear with *hΔx.* For the comparison of the two processes (embryonic vs. TADE), we take Δ*x* to be a fixed fraction of the apical cell width, w (Δ*x* = *C * w*). We note, that for a fixed cell volume, cells that are thin and wide consume more energy than cuboidal and adherent cells for the same contraction. However, TADE cells have only part of their volume in the tile, hence exhibit reduced thickness as well as reduced lateral dimensions.

Comparing the two cell geometries in this minimal toy model of horizontal springs, we get that the force required for contraction in TADE (taking h = 0.5um, w = 8um) compared to Drosophila embryo right before gastrulation (taking h = w = ~8um(30)) is times lower. This is a reduced order model and a full elastic simulation of the tissues, including all geometrical details, is required in future.

**Figure S1:**
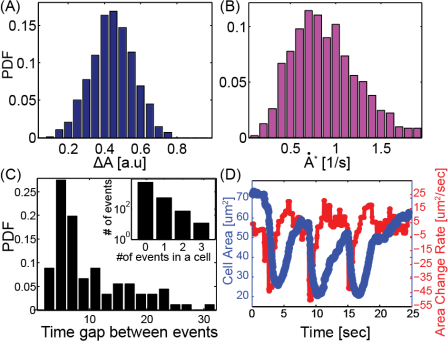
Additional statistical data. (A-B) PDFs of the amplitude and peak speed of the 746 contraction events from Fig. 2H-I, only in percentage units. (C) A PDF of the time interval between sequential contraction events. 86 such intervals were measured in 64 cells from the dataset in Fig 2. Inset: PDF of the number of events per cell in that dataset of 1 minute (D) Detailed example of a single cell that underwent sequential repetitive contractions (Mov. 6 inset). Blue is area, red is area change rate.

**Figure S2:**
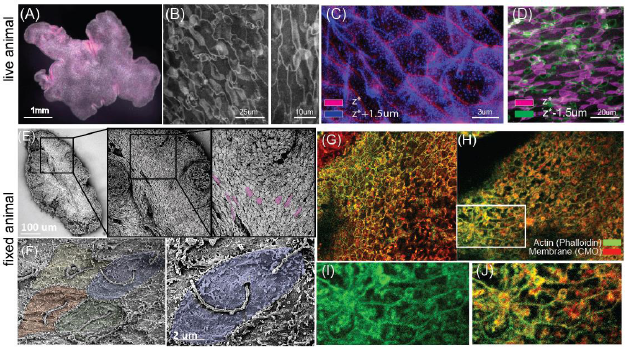
Cytoskeleton, cell and tissue morphology. (A) a dark field image of *T. adhaerens* from dorsal (top) view. (B-D) Fluorescence images from dorsal (top) view of the dorsal epithelium in vivo, using a live membrane stain (CMO) (B) Cell tiles are shown to stack/partially overlap (areas of higher intensity are due to more layers of fluorescent membrane). (C-D) Stack of two images in two colors representing two different planes of the same tissue segment (taken from a spinning disk confocal Z stack). In red is the plane showing tile-borders. In blue is the plane located 1.5 um towards outside of the animal, where membrane excursions are visible. In green is the plane located 1.5um inside the animal, where nucleus sacs are visible (E-F) SEM images of a fixed animal from dorsal (top) view. (E) A series of zoom-ins, from a whole animal to single cell shapes, some of which seem stretched as if responding to a global strain field. (F) Further zoom-in shows dorsal cells apical surface. A single cilia is at the center of each cell. The apical surface is ruffled. (G-J) Actin presence in dorsal cell tiles. (G) A confocal image of an animal that was relaxed before fixation and stained with CMO and phalloidin (methods). Actin filaments are seen only in cells’ periphery. (H) In an animal that did not undergo relaxation, the dorsal epithelium is due to some contraction prior to the fixation. Contracted cells are smaller and seen in the center of a ‘rosette’ (i.e. neighboring cells are expanded). (I-J) Zoom-in on a cells ‘rosette’. Filamentous actin signal is enhanced throughout the contracted cell tile.

**Figure S3:**
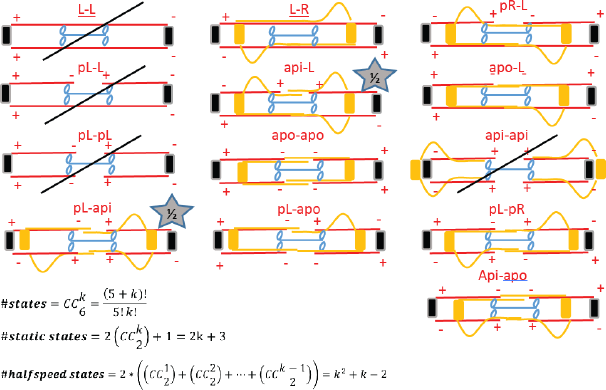
Possible microstates for a motor with 4 connected heads (k = 2). Depicted are all possible states of actin orientations relative to a motor filament with 4 connected heads. We exclude from the picture states that are symmetric to the depicted ones, either right-left or top-bottom. Therefore, instead of 21 possible states depicted are 13 states. The crossing line is marking static states that are not contracting at all. The star marks represents states that are contracting at half speed. The equations stated hold for any symmetric motor filament (even k).

## Movies S1-S10

1. Supplementary Video #1: Fluorescent fluctuations – the dorsal epithelium at low magnification. Dorsal (top) view of a live T. adhaerens using fluorescent membrane stain (CMO). The microscope stage is moving to keep the animal in the field of view, as it glides freely in the XY plane, in a 40um thick chamber. “Flashes” of increased fluorescence, about one second long, are seen sparsely across the tissue.
2. Supplementary Video #2: Sparse cellular contractions in TADE Dorsal (top) view of a live T. adhaerens using fluorescent membrane stain (CMO). The microscope stage is moving to keep the animal in the field of view, as it glides freely in the XY plane, in a 40um thick chamber. In addition, computational registration is performed to center the animal and correct for rotations. Individual cells are seen contracting in mostly uncorrelated locations and times. A contraction is correlated with an increased fluorescent signal. Right: On top of the original movie, color represents area change rate, as calculated via PIV. Red range marks expansions, blue range marks contractions. Low values, on both regimes, are excluded for clarity.
3. Supplementary Video #3: A radial contraction wave in TADE Dorsal (top) view of a live T. adhaerens using fluorescent membrane stain (CMO). The microscope stage is moving to keep the animal in the field of view, as it glides freely in the XY plane, in a 40um thick chamber. In addition, computational registration is performed to center the animal and correct for rotations. A radially propagating contraction wave is seen in the tissue. Right: On top of the original movie, color represents area change rate, as calculated via PIV. Red range marks expansions, blue range marks contractions. Low values, on both regimes, are excluded for clarity.
4. Supplementary Video #4: Uniaxial contraction waves in TADE Dorsal (top) view of a live T. adhaerens using fluorescent membrane stain (CMO). The microscope stage is moving to keep the animal in the field of view, as it glides freely in the XY plane, in a 40um thick chamber. In addition, computational registration is performed to center the animal and correct for rotations. Propagating contraction waves are seen in the tissue, mostly propagating in a uniaxial manner. Right: On top of the original movie, color represents area change rate, as calculated via PIV. Red range marks expansions, blue range marks contractions. Low values, on both regimes, are excluded for clarity.
5. Supplementary Video #5: Asynchronous contractile dynamics in high magnification and Individual cell cycles Dorsal (top) view of a live T. adhaerens using fluorescent membrane stain (CMO). The animal glides freely in the XY plane, in a 40um thick chamber. Contractions seem mostly asynchronous, yet the tissue is kept intact. A contraction is correlated with an increased fluorescent signal. Right: Individual cells are tracked computationally to show individual contraction cycles. Top right scale bar labells all single cell videos. The clock is joint to all parts of the video.
6. Supplementary Video #6: Sparse contractions mode in TADE – discrete and continuous analysis Dorsal (top) view of a live T. adhaerens using fluorescent membrane stain (CMO). The animal glides freely in the XY plane, in a 40um thick chamber. On top of the raw images, color represents area change rate, as calculated via PIV. Red range marks expansions, blue range marks contractions. Low values, on both regimes, are excluded. These thresholds on the continuous analysis follows individual cell borders accurately. Bottom right: discrete area measurements and statistical analysis is achieved by segmenting and tracking individual cells. The central cell at the inset is contracting three times. The clock is joint to all movie parts.
7. Supplementary Video #7: Hyperlinked contractions mode in TADE: Dorsal (top) view of a live T. adhaerens using fluorescent membrane stain (CMO). Majority of the cells are exhibiting repetitive active cycles of contraction/expansion. This state is related to high stress (animal experiences recent transfer/ sudden light increase /strong media flows).
8. Supplementary Video #8: Optical planes separation analysis Left: Dorsal (top) view movie of a live T. adhaerens using bright field illumination. The dorsal epithelium is at the focal plane, however as it is transparent only speckles on top of it are seen sharply. The ventral epithelium is out of the focal plane and is seen dark at the background. The layers are seen to move separately, sometimes in opposite directions. At time [00:07 sec] a contraction is seen at the top right of the animal. Middle: we split the movie by spatial frequency into two separate movies. The low and high frequency movies contain the ventral and dorsal data respectively. Top right: Planar displacement fields within each layer are calculated using PIV and represented as colored lines – dorsal (green) ventral (blue) and the difference between them (red). Bottom right: A heat map representing the local separation between the epithelia (the length of the difference vector). Values reach as high as 70um/sec (PIV is calculated on sec intervals).
9. Supplementary Video #9: Ionomycine effect on TADE Dorsal (top) view of a live T. adhaerens using fluorescent membrane stain (CMO). At time [00:04] the drug Ionomycine is introduced. All cells immediately respond in contraction. Minimal area is achieved within approximately 10sec. The contracted state lasted until drug was washed away.
10. Supplementary Video #10: TADE cell shape variability under spontaneously generated stress Dorsal (top) view movies of a live T. adhaerens using fluorescent membrane stain (CMO). The movies demonstrate that TADE cells experience extreme fluctuations in the stress levels and their orientations at the scale of seconds. These fluctuations originate from the animal locomotion (the ciliary array at the ventral side gliding on the surface) and the overall animal shape changes. At the same time scales, cells change their size and their shape, and accommodate a continuous, intact tissue at all times.

